# GFAP: ultra-fast and accurate gene functional annotation software for plants

**DOI:** 10.1101/2022.01.05.475154

**Authors:** Dong Xu, Kangming Jin, Heling Jiang, Desheng Gong, Jinbao Yang, Wenjuan Yu, Yingxue Yang, Jihong Li, Weihua Pan

## Abstract

Sequence alignment is the basis of gene functional annotation for unknow sequences. Selecting closely related species as the reference species should be an effective way to improve the accuracy of gene annotation for plants, compared with only based on one or some model plants. Therefore, limited species number in previous software or website is disadvantageous for plant gene annotation.

Here, we collected the protein sequences of 236 plant species with known genomic information from 63 families. After that, these sequences were annotated by pfam, Gene Ontology (GO) and Kyoto Encyclopedia of Genes and Genomes (KEGG) databases to construct our databases. Furthermore, we developed the software, **G**ene **A**nnotation **S**oftware for **P**lants (GFAP), to perform gene annotation using our databases. GFAP, an open-source software running on Windows and MacOS systems, is an efficient and network independent tool. GFAP can search the protein domain, GO and KEGG information for 43000 genes within 4 minutes. In addition, GFAP can also perform the sequence alignment, statistical analysis and drawing. The website of https://gitee.com/simon198912167815/gfap-database provides the software, databases, testing data and video tutorials for users.

GFAP contained large amount of plant-species information. We believe that it will become a powerful tool in gene annotation using closely related species for phytologists.

## Introduction

Gene functional annotation (GFA) is an important part for bioinformatic analysis, such as genomics (Cheng et al., 2021), transcriptome (Fernandez-Valverde et al., 2015) and gene family analysis (Martin et al., 2010). Furthermore, GFA can also provide vital guidance for wet-lab biologists to explain the life phenomenon (Wei et al., 2017). Therefore, GFA plays important roles in almost every aspect of plant studies.

However, annotation errors were continuously reported in various studies (Jones et al., 2007; Bayer et al., 2018). The absence of befitting species models is an essential factor for these annotation errors (Kim et al., 2020). For example, *Arabidopsis thaliana*, a common model plant in many GFA websites, has significant differences in xylem development of woody plants (Jiao et al., 2012; Bu et al., 2021). Selecting *Arabidopsis thaliana* as the reference species to annotate the xylem developmental genes of woody plants may result in annotation errors. To date, although more and more plant genomes have been sequenced, available plant models for gene annotation were still scarce. For example, only two species were plants (*Arabidopsis* and rice) among the 15 species collected in GeneCodis (Nogales-Cadenas et al., 2009). The similar phenomenon can also be found in other websites (Medina et al., 2010; Kuleshov et al., 2016). Large amount of unknow genes were discovered and remained to be annotated, with the development of high-throughput sequencing technology. Therefore, it is higher requirements in annotation efficiency of gene annotation software. In other word, an ideal annotation tools should not only contain large amount of plant models but also have a high efficiency in gene annotation.

To solve the problems of plant-model absence, the genes of total 200 plant species from 60 families were annotated by the database of the protein families database (pfam), Gene Ontology (GO) and Kyoto Encyclopedia of Genes and Genomes (KEGG) (Kanehisa and Goto, 2000; Consortium, 2004; Finn et al., 2014). The annotated results were added into our databases. To improve annotation efficiency, we optimized our databases structures and developed a novel software, **G**ene **A**nnotation **S**oftware for **P**lants (GFAP), for calling relevant information from our databases to complete the annotation process. For example, in testing phase, 43000 genes can be annotated by their protein domains within 56 seconds (Video S1). Furthermore, there was no limit of GFAP in the size of input files, which can be essential for dealing with big biological data. Therefore, we believe that GFAP could be a useful tool for phytologists to solve the problems in GFA of plants.

## Results

### Database information

The basic information of our databases was listed in the Table S1. Total 125 species were concluded into our databases, ranging from algae, mosses, gymnosperms to monocots and dicotyledon. Among of them, the data of 51 species from 51 families were added into GFAP. Other data can be freely downloaded from the website of https://gitee.com/simon198912167815/gafp-database, and can be utilized by GFAP when added into the indicated folders. Figure 1 showed the database-construction process and the roles of databases (including DNA/protein-sequences, protein-domain, GO and KEGG databases) in using GFAP.

**Figure 1.**
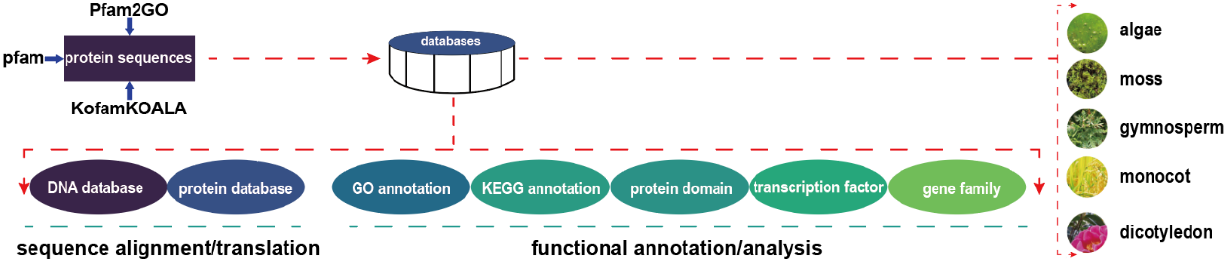
The construction of GFAP databases and the contained data. All protein domains in full sequences were detected, and annotated the corresponding information. The GFAP databases were thus constructed. These protein sequences were gained from five categories of plants, ranging from algae, moss, gymnosperm to monocot and dicotyledon. Five kinds of data were contained in the databases. In detail, DNA and protein data were utilized for sequence alignment. The data of GO, KEGG and protein domain were used for the annotating process.

### Overview of the functions and workflow of GFAP

Four modules were constructed in GFAP (Figure 2a). The Alignment module was responsible for sequence alignment. Users can align their sequences to a selected database. For example, if obtaining some sequences from Crassulaceae plants, the users can select *Kalanchoe fedtschenkoi* (a Crassulaceae plant in database) for alignment. The alignment process was completed by the diamond software (Hernández-Salmerón and Moreno-Hagelsieb, 2020) to increase the alignment efficiency. Here, we strongly encouraged users to utilize the protein sequences for alignment. However, considering that some users may not have the protein sequences, GFAP still supported the DNA alignment, and users need to download the relevant DNA databases from our website. After that, users can flexibly choose the types of functional annotation (Figure 2b). The protein domains can be detected using the superHMM module (Figure 2a), and the results can help users predict gene functions or identify the members of their interested gene families. The annotation results of GO and KEGG databases can be obtained from the GO analysis and KEGG analysis module, respectively. Furthermore, the processes of statistical analysis and drawing can also be completed by the relevant functions of GFAP (Figure 2a).

**Figure 2.**
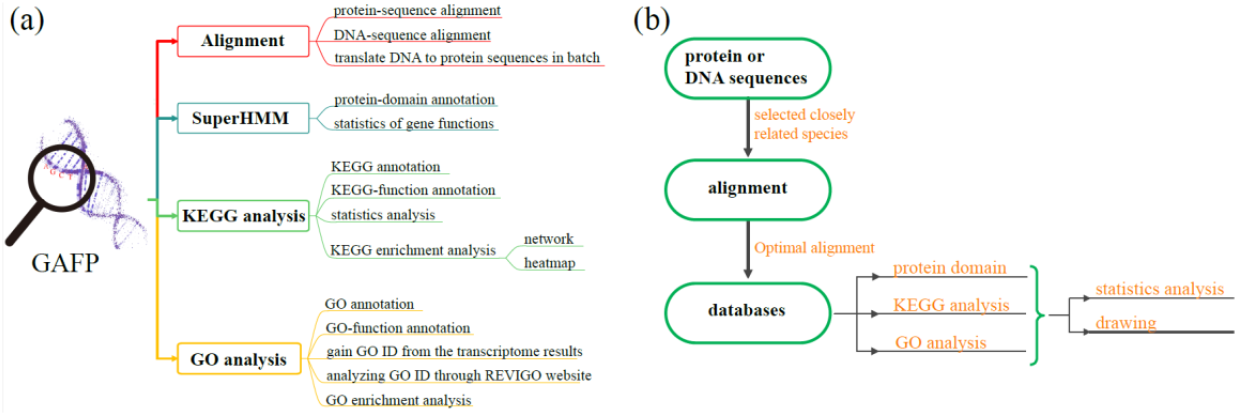
Functions and workflow of GFAP. (a) overview of GFAP functions. Total four modules (including sequence alignment, protein-domain query, KEGG ID and GO ID) were contained in GFAP. They are responsible for the sequence alignment, translation, identification of protein domains, KEGG annotation, GO annotation, statistics analysis and drawing. (b) the workflow of GFAP. Users can annotate their sequences in just four steps.

### The characters of GFAP

Compared with previous websites or software (Table S2), GFAP has the following characteristics:

1. GFAP was specially designed for gene annotation of plants. Our databases contained the annotated information of over 200 plant species, which can allow users freely select the closely related species to annotate their interested genes.
2. Highly efficient gene annotation. The optimized data structure and efficient extracted tools can annotate thousands of genes within several seconds or minutes (Video S1, in Windows 10, the random access memory was 8 GB).
3. More comprehensive annotation information. The GFAP databases contained the information of all protein domains of genes. Therefore, the annotation results of GFAP can provide more information for users, which can help phytologists better understand their interested genes. For example, The domains related with phosphatase and nucleotidase can be simultaneously detected in the sequences of Bradi2g62150.2.p (Table 1), indicating that Bradi2g62150.2.p may act in the process relevant with phosphatase and nucleotidase.
4. Systematic analysis. In addition to the functional annotation, statistical analysis and drawing can also be performed using GFAP Windows version. For example, users can make GO clustering analysis utilizing the “GO analysis” module (Figure 3 a). The heatmap and network can also be drawn by the “KEGG analysis” (Figure 3b and c). Furthermore, the format of GFAP-output files can meet with the requirements of other websites or software, such as REVIGO (http://revigo.irb.hr/) and clusterProfiler (Yu et al., 2012).
5. Fewer limits in using GFAP. The functions of GFAP can be completed by point-and-click icons instead of inputting any command lines. The prompt information on the GFAP surface (Figure 3d) can help users run the GFAP without any barriers. GFAP can perform its functions independent with internet. Furthermore, there were not limits of GFAP to the input-file size. For example, the protein-domain annotation for a 91.1 Mb protein-sequence file can be completed within three minutes (Video S2).

**Figure 3.**
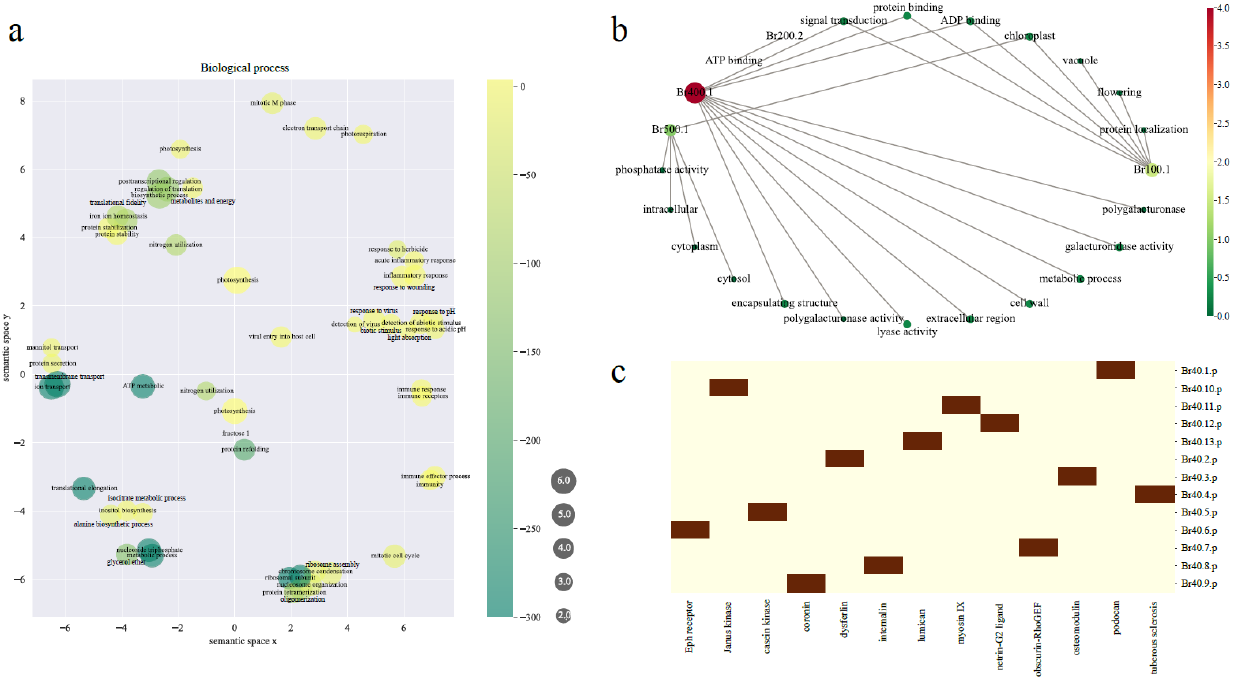
Drawing functions and interface features of GFAP. Enrichment analysis of GO can be completed by GFAP, with the help of REVIGO (a). The network and heatmap can also be finished using GFAP (b and c). The prompt information on the interface of GFAP helped users run GFAP accurately (d).

**Table 1.**
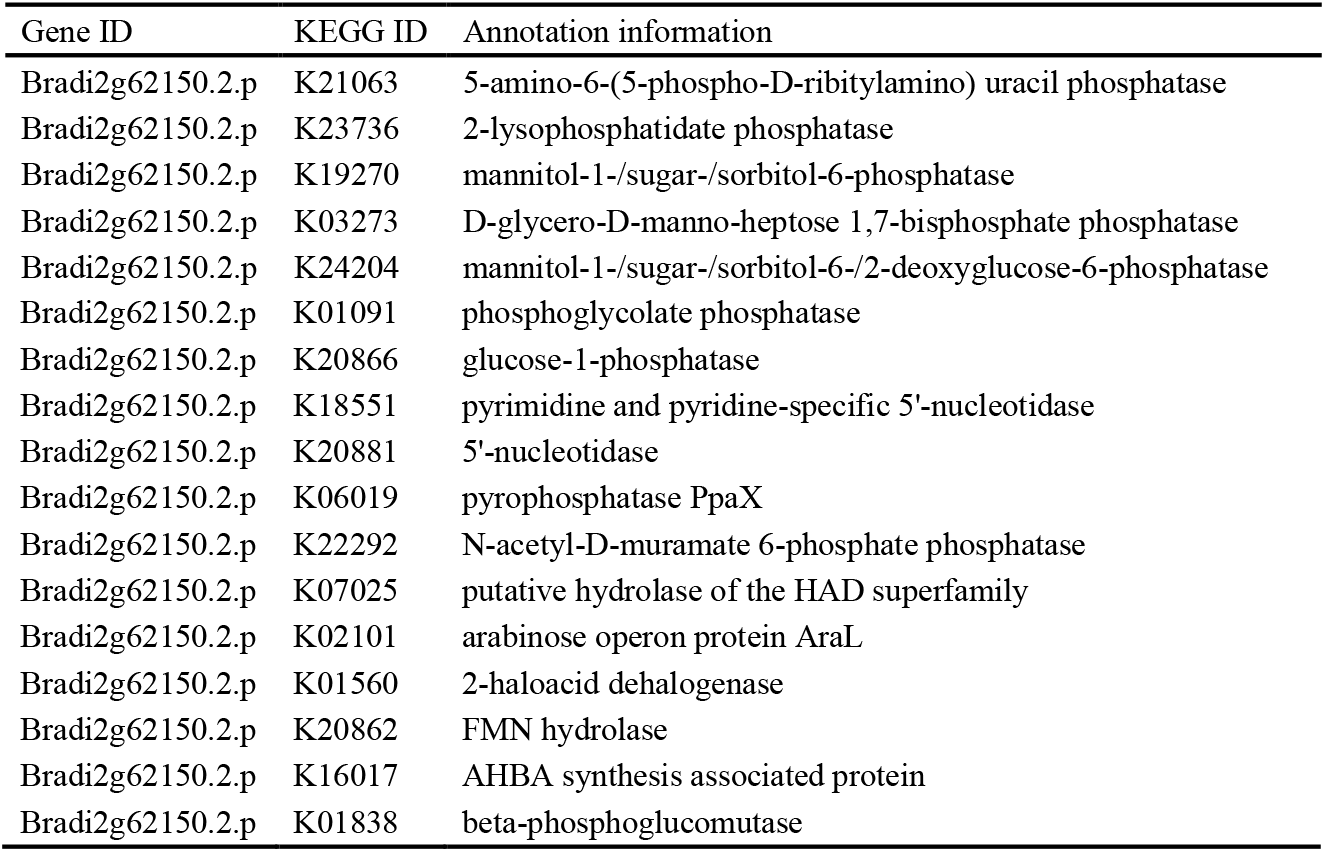
The annotation information for Bradi2g62150.2.p.

| Gene ID | KEGG ID | Annotation information |
| --- | --- | --- |
| Bradi2g62150.2.p | K21063 | 5-amino-6-(5-phospho-D-ribitylamino) uracil phosphatase |
| Bradi2g62150.2.p | K23736 | 2-lysophosphatidate phosphatase |
| Bradi2g62150.2.p | K19270 | mannitol-1-/sugar-/sorbitol-6-phosphatase |
| Bradi2g62150.2.p | K03273 | D-glycero-D-manno-heptose 1,7-bisphosphate phosphatase |
| Bradi2g62150.2.p | K24204 | mannitol-1-/sugar-/sorbitol-6-/2-deoxyglucose-6-phosphatase |
| Bradi2g62150.2.p | K01091 | phosphoglycolate phosphatase |
| Bradi2g62150.2.p | K20866 | glucose-1-phosphatase |
| Bradi2g62150.2.p | K18551 | pyrimidine and pyridine-specific 5'-nucleotidase |
| Bradi2g62150.2.p | K20881 | 5'-nucleotidase |
| Bradi2g62150.2.p | K06019 | pyrophosphatase PpaX |
| Bradi2g62150.2.p | K22292 | N-acetyl-D-muramate 6-phosphate phosphatase |
| Bradi2g62150.2.p | K07025 | putative hydrolase of the HAD superfamily |
| Bradi2g62150.2.p | K02101 | arabinose operon protein AraL |
| Bradi2g62150.2.p | K01560 | 2-haloacid dehalogenase |
| Bradi2g62150.2.p | K20862 | FMN hydrolase |
| Bradi2g62150.2.p | K16017 | AHBA synthesis associated protein |
| Bradi2g62150.2.p | K01838 | beta-phosphoglucomutase |

### The accuracy of GFAP

The accuracy of GFA is an important issue for users. Two aspects of comparisons were made to demonstrate the GFAP accuracy. The comparison of annotated results should be made among different databases of GFAP firstly, as there should be some similarities in annotation results among different databases (even though they described different aspects of a gene). The protein-sequence file of *Brachypodium distachyon* was downloaded from the phytozome website, and we randomly selected 500 genes for annotation. As *B. distachyon* is in *Gramineae* family, we chose *Oryza sativa* as the reference species. The results were listed in Table S3, and the similar results of the three databases were marked by red color. Figure 4 showed the statistical result. Total 372, 347 and 410 genes were annotated by the protein-domain, KEGG and GO databases, respectively. Among of them, the annotations of 414 genes obtained from at least two databases were similar with each other. The functions of 44 genes remained to be unknown. These results indicated that the annotation results of GFAP had high accuracy, as a large amount of genes were annotated by similar information from different databases.

**Figure 4.**
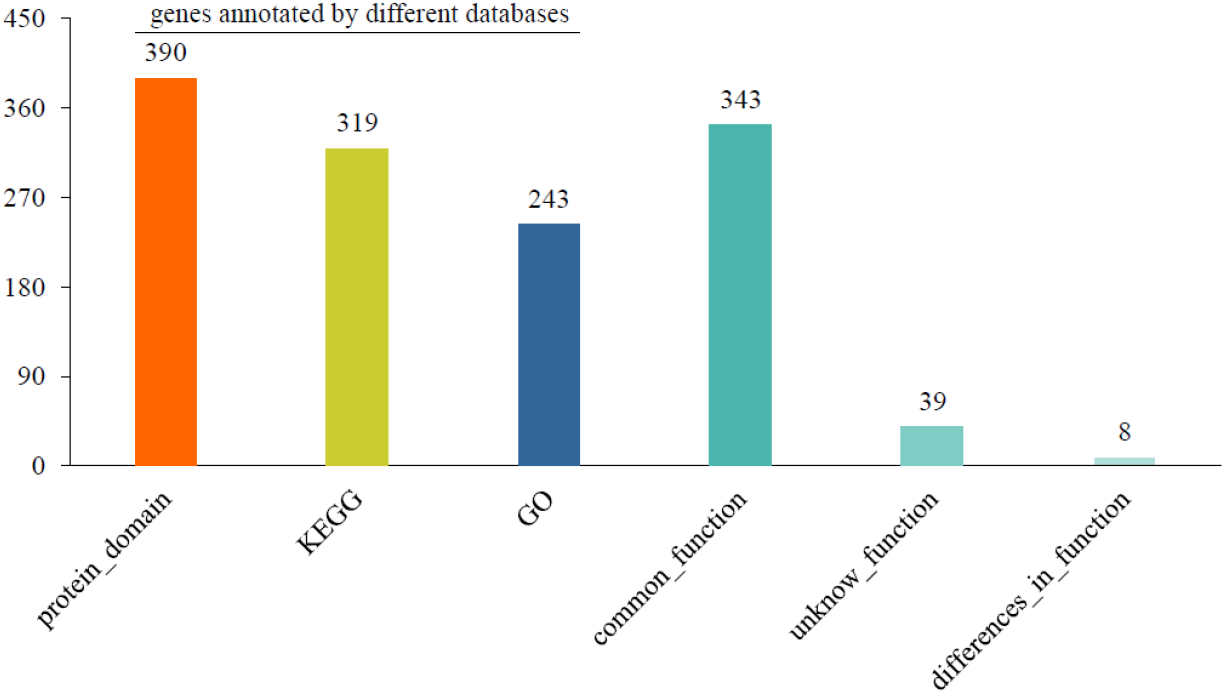
The number of annotated genes with different reference species.

Secondly, we compared the annotation results of GFAP with the functions demonstrated by wet-lab biologists (Table 2). We randomly selected ten genes in poplars, and chose *Populus deltoides* as the reference species for gene annotation. The annotated results were listed in Table S4, and the partial results were showed in Table 2. We found that the annotation results were highly consistent with the published functions. For example, the *NatA* was a N-terminal acetyltransferase (Zhu et al., 2014). The acetyltransferase domain was detected in the “superHMM” module of GFAP. Similarly, the N-acetyltransferase activity was found using the functions of GO annotation. While, the sequence was directly defined as the N-acetyltransferase by KEGG annotation. In summary, based on the above results, we believed that the annotation results of GFAP were highly accurate.

**Table 2.** Comparison of the annotated information by GFAP with the published functions of genes

| Gene ID | Publication information | Functions in articles | Protein-domain (GFAP) | GO annotation (GFAP) | KEGG annotation (GFAP) |
| --- | --- | --- | --- | --- | --- |
| *LysoPL2* | (Jiang et al., 2021) | Lysophospholipase | Phospholipase/carboxylesterase | Lysophospholipase activity | Lysophospholipase |
| *NatA* | (Zhu et al., 2014) | N-terminal acetyltransferase | Acetyltransferase domain | N-acetyltransferase activity | N-acetyltransferase |
| *PdERECTA* | (Xing et al., 2011) | Leucine-rich repeat<br>receptor-like kinase | Leucine rich repeat kinase domain | Protein kinase activity | Leucine-rich repeat domain-containing<br>protein |
| *Pop14A9* | (Lafarguette et al., 2004) | Fasciclin-like<br>arabinogalactan | Fasciclin domain | Plant-type secondary cell wall<br>biogenesis | Fasciclin 1 |
| *PtHMA4* | (Adams et al., 2011) | Heavy metal regulation | Heavy-metal-associated domain | Metal ion transport | Cd<sup>2+</sup>-exporting ATPase |
| *PtaZFP2* | (Martin et al., 2014) | Zinc finger protein | C2H2-type zinc finger | Response to wounding | Zinc finger protein |
| *PtC3H17* | (Chai et al., 2014) | CCCH zinc finger protein | Zinc finger C-x8-C-x5-C-x3-H type | Metal ion binding | RING /CCCH-type zinc finger protein |
| *PttAMT1.2* | (Selle et al., 2005) | Ammonium importer | Ammonium transporter family | Ammonium transport | Ammonium transporter, Amt family |
| *PttSLAH3* | (Jaborsky et al., 2016) | Anion channel | Voltage-dependent anion channel | Anion channel activity | Tellurite resistance protein |
| *Ptxerico* | (Kim et al., 2020) | RING-H2 Zinc Finger | RING-H2 zinc finger domain | Zinc ion/ protein binding | Zinc finger -like/RING finger protein |

## Discussion

GFA plays important roles in many aspects of plant studies. For example, GFA can help wet-lab phytologists explain the developmental process of plants (Wei et al., 2017). GFA is the basis for gene family analysis (Xiao et al., 2018). In addition, it is also an important part for genomics and transcriptomics (Bengtsson et al., 2014; Chen et al., 2021). Therefore, the results of GFA will profoundly influence the development of plant science. Compared with previous websites (Huang et al., 2009; Zhou et al., 2019), we constructed GFA databases for 124 plant species, and developed GASF software to efficiently annotate the unknow-function genes. Over 52 families of plants were concluded into our databases. We believed that GASF can annotate most plant species lacking of genomic information with the information of closely related species.

Sequence alignment is the basis for GFA (Xiao et al., 2018). For this reason, the results of GFA are highly affected by the similarities between sequences in databases and the function-unknow sequences. This may also be an important cause of GFA deviation. In this study, we found that compared with choosing closely related species, 30-40 genes from *B. distachyon* were not annotated when selecting *A. thaliana* as the reference species (Figure 4). This result further highlights the importance of selecting closely related species for GFA, as high sequence similarities can be found between closely related species. Meanwhile, in GASF, the diamond program was chosen for sequence alignment. The accuracy of diamond is higher than that of the traditional blast program (Hernández-Salmerón and Moreno-Hagelsieb, 2020). The above factors guaranteed the accuracy of the GASF annotation results.

## Materials and Methods

In this study, the protein and DNA sequences were downloaded from the website of phytozome (https://phytozome-next.jgi.doe.gov/), TPIA (http://tpia.teaplant.org/index.html), eplant (http://eplant.njau.edu.cn/), EnsemblPlants (http://plants.ensembl.org/info/data/ftp/index.html) and NCBI (ftp://ftp.ncbi.nlm.nih.gov/genomes/refseq/plant). After that, the longest transcript of each gene was remained using the functions of SPDE (Xu et al., 2021).

### Protein-domain database

The database of Pfam-A (Finn et al., 2013) was used for identification of protein domains in batch, with the help of hmmpress and hmmscan programs (Malhotra and Sowdhamini, 2013). To accelerate the speed of annotations and reduce the size of the database, the gene ID and its protein-domain name were remained, and other information was removed from the annotation results.

### KEGG database

KEGG database was built by KofamKOALA (Aramaki et al., 2019) using protein sequences. As stated above, only gene ID and the relevant KEGG ID were remained.

### GO database

After obtaining the Pfam ID for each gene, the database of Pfam2GO (https://rdrr.io/github/missuse/ragp/man/pfam2go.html) was used to identify the GO ID for genes (Mitchell et al., 2015). Furthermore, the software of blast2go (Conesa et al., 2005) was also utilized for confirming these results.

### The development of GFAP

GFAP was developed by python language using 3.8 version.

### Conclusion

GFAP is a highly efficient and accurate tool for gene functional annotation. Its accuracy indicated that it can play important roles in predicting the gene functions for wet-lab phytologists. Moreover, the high efficiency and accuracy revealed that it can be used for functional genomics and transcriptome analysis. At the same time, lots of plant-species information can be available for related species annotation, and this process provided a new annotated method for species with unknow genomic information.

## Acknowledgements

This study was supported by the Zhejiang Science and Technology Major Program on Agricultural New Variety Breeding (2021C02070-1), the National Natural Science Foundation of China (31872168).

